# EYA1/EYA2 and EYA3/EYA4 act as stage-specific SIX cofactors in embryonic and adult regenerative skeletal myogenesis

**DOI:** 10.64898/2026.05.20.726470

**Authors:** Camille Viaut, Maud Wurmser, Edgar Jauliac, Laura Ben Driss, Stéphanie Backer, Rouba Madani, Fayez Issa, Iryna Pirozkhova, Athanassia Sotiropoulos, Helge Amthor, Pascal Maire

## Abstract

*Eya3* and *Eya4* are two *Eya* genes expressed in adult myogenic stem cells, where they may act as SIX cofactors. We analyzed muscle regeneration in single and compound *Eya3* and satellite cell-specific *Eya4* mutant mice. A kinetic analysis of muscle regeneration after Notexin injury of the Tibialis Anterior revealed no major phenotype at 4, 14, and 30 days after injury in terms of PAX7+ cell number and myofiber cross-sectional area in *Eya3* mutants, while all parameters were decreased in *Eya4* mutants and further worsened in *Eya3/Eya4* double mutants, in which we also observed a modification of the myofiber phenotype at 30 days after injury. Satellite cells were cultured ex vivo and *Eya4* deletion was induced by Ad-Cre-mediated recombination. While single *Eya3* mutant cells showed normal proliferation and differentiation, double mutant cells exhibited normal proliferation but failed to fuse. Analysis of their transcriptome revealed that the expression of *Myomixer*, *Follistatin*, and *Noggin* was severely downregulated specifically in double mutant cells, explaining their fusion deficiency. To gain a better understanding of the involvement of *Eya* genes during embryonic development and the genesis of PAX7+ myogenic stem cells, we analyzed *Eya1 / ;Eya2 / , Eya3 / , Eya4 /* , and *Eya3 / ;Eya4 /* E18.5 mutant fetuses at the limb and craniofacial levels. In *Eya1 / ;Eya2 /* fetuses, we confirmed the absence of distal limb muscles and observed reduced craniofacial muscles. In *Eya3 / ;Eya4 /* fetuses, craniofacial myogenesis appeared preserved and PAX7+ myogenic stem cells were present.

**Background:** The *Eyes absent* (*Eya*) genes encode transcriptional co-activators and phosphatases that function within the PAX-SIX-EYA-DACH (PSED) regulatory network. In skeletal muscle, EYA proteins cooperate with SIX homeoproteins to control myogenic gene expression during both embryonic development and adult regeneration. While *Eya1* and *Eya2* are predominantly expressed in embryonic myogenic progenitors and *Eya3* and *Eya4* are the dominant paralogs in adult satellite cells (SC), the specific and redundant contributions of individual family members to myogenesis remain poorly characterized.

**Methods:** We analyzed compound *Eya* mutant mice during adult *Tibialis anterior* muscle regeneration and during embryogenesis. We complemented this analysis by performing *ex vivo* myogenic stem cell cultures from compound *Eya* mutants and examining their fusion capacity.

**Results:** Analysis of muscle regeneration following Notexin injury revealed that *Eya2* and *Eya3* single mutants display no major regenerative deficit. In contrast, satellite cell-specific deletion of *Eya4* (*Eya4^sc/sc^*) caused a transient impairment of early regeneration, with reduced numbers of smaller regenerating MYH3+ (embryonic myosin heavy chain) myofibers and a transient decrease in SC number at 4 days post-injury (dpi). Compound *Eya3^-/-^;Eya4^sc/sc^*double mutants showed a more severe and persistent phenotype, with decreased myofiber cross-sectional area, reduced myonuclear accretion, accumulation of PAX7+ cells associated with regenerated myofibers, and altered fiber-type composition at 14 and 30 dpi. *Ex vivo* analysis of double mutant SCs revealed a specific and complete blockade of myogenic fusion without defects in proliferation or MYOD expression. Transcriptomic analysis identified severe downregulation of *Myomixer*, *Noggin*, and *Follistatin* in differentiating *Eya3^-/-^;Eya4^-/-^* SCs. Open-access SIX1 and SIX4 ChIP-seq publicly available data confirmed direct binding at the *Myomixer*, *Noggin*, and *Follistatin* loci, supporting a direct SIX-EYA transcriptional mechanism. In parallel, embryonic analysis demonstrated that *Eya1^-/-^;Eya2^−/−^*E18.5 fetuses lack distal limb musculature and display severe craniofacial muscle hypoplasia, while in *Eya3^-/-^;Eya4^−/−^*fetuses limb and craniofacial musculature developed with no detectable defects.

**Conclusions:** These results reveal distinct temporal requirements for EYA proteins in skeletal muscle: EYA1 and EYA2 are essential SIX cofactors for embryonic myogenic fate acquisition in hypaxial and craniofacial progenitors, while EYA3 and EYA4 act redundantly in adult satellite cells to enable myogenic fusion by maintaining BMP antagonist expression and *Myomixer* activation downstream of the SIX-EYA transcriptional complex.

## Background

During embryogenesis, skeletal muscle formation is initiated in distinct embryonic territories, and myogenic fate acquisition is triggered by the expression of one of the Myogenic Regulatory Factors (MRFs) of the *MyoD* family [1,2]. The absence of MRFs precludes the genesis of embryonic myogenic cells [3], while the loss of *Myf5* and *MyoD* in adult myogenic stem cells impairs adult muscle regeneration [4].

SIX and EYA proteins participate in the activation of MRFs during embryonic development [5–7], and also during adult muscle regeneration [8–11]. In addition, SIX homeoproteins act through feedforward regulatory mechanisms and are required at all steps of muscle development [12].

During development *Eya* genes are expressed in several tissues in association with *Six* genes and control the morphogenesis of kidney, ear, thymus, cranial ganglia, sensory placodes and skeletal muscles [5,13–15]. *Eya2^-/-^* and *Eya3^-/-^* mutant mice are viable and fertile, with *Eya3^-/-^*mice being smaller in size [6,16], while *Eya1^-/-^*mice die at birth due to kidney hypoplasia [14]. *Eya4^-/-^* mice are smaller in size, have severe hearing deficits; in addition, the males are sterile [17]. While expression of *Six* and *Eya* genes are downregulated in most adult tissues they remain expressed in post-mitotic adult myofibers as well as their associated myogenic stem cells [18].

Myogenic stem cells acquire a satellite cell position at the end of fetal development. Once localized between the myofiber and the basal lamina, they contribute to myofiber growth and regeneration [2,19–22]. This homing process is compromised in *Six1Six4* mutant mice [21]. Adult muscle regeneration is initiated following myofiber breakdown through activation of PAX7+ myogenic stem cells, called satellite cells (SCs), which are essential for muscle regeneration [23–26] and hypertrophy [27–31]. Although *Six1* and *Six4* are required for efficient adult muscle regeneration, the specific roles of their associated EYA cofactors during embryonic and adult myogenesis remain poorly understood.

In the present study we refine the role of *Eya* paralogs during both adult muscle regeneration and embryonic development. We show that all four *Eya* genes are expressed in myogenic stem cells in the adult and in the embryo. Analysis of single and double *Eya* mutant revealed the important role of *Eya4* in myogenic stem cells during adult muscle regeneration and in *Eya1* and *Eya2* for limb and craniofacial muscle development.

## Materials & Methods

### Mice and animal care

Animal experimentation was carried out in strict accordance with the European convention STE 123 and the French national charter on the Ethics of Animal Experimentation. Protocols were approved by the Ethical Committee of Animal Experiments of Institut Cochin, CNRS UMR 8104, INSERM U1016, and by the Ministère de l’éducation nationale, de l’enseignement et de la recherche, no. APAFIS#15699-2018021516569195. Mice were maintained at 22 ± 2°C, with 30–70% humidity and a 12h/12h dark/light cycle. Wild-type (WT) C57BL/6N mice were purchased from Janvier Laboratories. All mutant mice used in this project were 2-to 6-month-old and backcrossed onto a C57BL/6N background.

The *Eya4^flox^* allele was generated from the *Eya4tm1a(KOMP)Wtsi* mouse strain, provided by the KOMP Repository (www.komp.org), by crossing with transgenic mice expressing a ubiquitous Flipase to delete the PGK-neo selection cassette flanked by FRT sites, yielding *Eya4^flox/+^* F1 progeny. This allele carries loxP sites flanking exons 12 and 13 of *Eya4* (Fig. S2B). *Eya4^flox/+^*mice were crossed with *EIIaCre* animals expressing Cre recombinase ubiquitously under the E2A promoter [32] to generate the *Eya4^-/+^* strain. The *Eya3-lacZ* knock-in (*Eya3^-/+^*) mice were obtained from Söker et al. [16]. *Eya3^-/-^;Eya4^-/+^*animals were crossed to obtain *Eya3-/-;Eya4-/-* and *Eya3^-/-^;Eya4^+/+^*fetuses, and *Eya4^-/+^* animals were intercrossed to obtain *Eya4^-/-^*fetuses.

Satellite cell (SC)-specific *Eya4* conditional mutant mice (*Eya4^flox/flox^;Pax7^CreERT2/+^*) were generated by crossing *Eya4^flox/flox^* animals with *Pax7^CreERT2/+^*animals [24], in which a tamoxifen-inducible CreERT2 transgene is placed at the endogenous *Pax7* locus. *Eya3^-/-^ ::Eya4^flox/flox^::Pax7^CreERT2/+^* mice were obtained by further crossing with *Eya3^-/+^* animals. To generate a constitutive *Eya3^-/-^;Eya4^-/-^* line, *Eya3^-/-^;Eya4^flox/flox^;Pax7^CreERT2/+^* mice were crossed with *EIIa-Cre* mice, producing *Eya3^-/+^;Eya4^-/+^* mice that were subsequently intercrossed.

*Eya1^-/-^*, *Eya2^-/-^*, *Eya3^-/-^*, *Eya4^-/-^*, *Eya1^-/-^ ;Eya2^-/-^*, and *Eya3^-/-^;Eya4^-/-^*E18.5 mutant fetuses, along with their littermate controls, were obtained by intercrossing *Eya1^-/+^ ;Eya2^-/+^*mice [6] and *Eya3^-/+^ ;Eya4^-/+^* mice (this work), respectively. Genotyping of mice and embryos was performed using specifically designed primers (Table S1).

For conditional *Eya4* deletion in satellite cells, tamoxifen was administered intraperitoneally to 2–6 month-old mice daily for 5 consecutive days prior to muscle injury. Tibialis anterior (TA) injury was performed on anesthetized animals by a single intramuscular injection in the *Tibialis anterior* of 20 µl of Notexin (5 µg/ml NTX); mice were allowed to recover for 4 to 30 days post-injury. Mice were euthanized 4, 14 or 30 days after Notexin injury. Muscles were freshly frozen in O.C.T. Embedding Matrix compound (Cellpath), snap frozen in isopentane cooled in liquid nitrogen, and stored at -80°C until use. Transverse *Tibialis anterior* sections (10 µm in thickness) were obtained using a Leica cryostat and collected on SuperFrost-Plus slides (ThermoFisher).

### Immunohistochemistry on muscle sections

Adult *Tibialis anterior* muscle cryosections were fixed in 4% PFA for 20 min at RT and permeabilized with cooled-methanol for 6 min. Antigen retrieval was performed with the Antigen Unmasking Solution (pH 6, Vector H-3300) at 95°C for 10 min, and cooled on ice for 20 min. Sections were washed in 1X PBS and blocked with 5% goat serum (GS)/0.5% Triton/0.1M glycine/4% BSA/1X PBS for at least 1 hour. Sections were incubated with primary antibodies overnight at 4°C. After 1-hour incubation at RT with secondary antibodies and Hoechst (nuclei staining), slides were mounted in Mowiol mounting medium and kept at 4°C until image acquisition. Primary and secondary antibodies are listed in (Table S2). For immunostaining against MYH4, MYH2 and MYH7, cryosections were used without fixation. They were rehydrated in 1X PBS, directly blocked with 10% goat serum/1X PBS for 30 min at RT, and incubated with antibodies as previously described.

### Fetuses preparation

Fetuses were staged by taking the appearance of the vaginal plug at embryonic day 0.5 (E0.5). They were harvested 18.5 days post-fertilization, decapitated and their skin was removed. Isolated head, trunk and limbs (fore-and hindlimbs from one side) were fixed in 4% paraformaldehyde (PFA) for 30 min at room temperature (RT) and kept in 15% then 30% sucrose-PBS at 4°C overnight. Head, trunk and limbs were embedded into O.C.T. Embedding Matrix compound (Cellpath), snap frozen in isopentane cooled on dry ice, and stored at - 80°C. Transverse sections (10 µm in thickness) at eye, heart and limb levels were collected on SuperFrost-Plus slides (ThermoFisher). snRNA-seq from Notexin induced TA regeneration will be detailed elsewhere (Viaut et al, unpublished).

### Immunohistochemistry on fetus sections

Fetus sections were rehydrated in 1X PBS. For most immunostainings, antigen retrieval was performed in citrate buffer (pH 6; Vector H-3300) at 95°C for 10–15 min, followed by 20 min of cooling. Sections were then permeabilized in 0.5% Triton/0.1M glycine/1X PBS for 30 min and blocked with 10% horse serum/0.5% Triton/0.1M glycine/1.5% BSA/1X PBS for at least 1–3h at RT. Sections were incubated with primary antibodies overnight at 4°C, then with secondary antibodies and Hoechst (nuclei staining) for 45 min–1h at RT; all antibodies were diluted in blocking solution. Primary and secondary antibodies are listed in Table S2.

For PAX7 immunostaining, antigen retrieval was performed as described above, after which sections were additionally permeabilized with cold −20°C acetone for 10 min and air-dried for 10 min before blocking and antibody incubations. For CD31 immunostaining, sections were permeabilized with cold −20°C acetone for 10 min and air-dried for 10 min, then blocked and incubated with antibodies as described above (no prior antigen retrieval).

For SIX1 and EYA4 immunostainings, a tyramide signal amplification (TSA) step was required. After incubation with a biotinylated secondary antibody, sections were incubated with horseradish peroxidase (HRP)-conjugated streptavidin for 30 min, then revealed with either Alexa Fluor 488 or Alexa Fluor 546 Tyramide Reagent (SuperBoost TSA kit, Thermo Fisher Scientific; or Invitrogen #B40954) in 30% H O /amplification buffer (PerkinElmer) for 10 min.

Immunostained sections were mounted in Mowiol mounting medium and kept at 4°C until image acquisition.

### Fluorescence Activated Cell Sorting (FACS)-mediated satellite cell isolation

Skeletal muscle-derived primary myoblasts were obtained from hindlimb muscles of WT and *Eya3^-/-^Eya4^flox/flox^* mice after enzymatic digestion followed by FACS purification. Briefly, fore- and hindlimbs were dissected out, finely minced to a pulp and digested at 37°C for 2 hours in Digestion Solution (Dispase II (3mg/mL), collagenase A (100 µg/µL), 0.4 mM CaCl2, DNase I (10 µg/mL), 5 mM MgCl2 in 1X HBSS (Roche and Life Technologies)). Muscle extracts were washed with 1X HBSS/0.2% BSA and filtered through 100-µm and 40-µm strainers (Corning Life Science). Erythrocytes were eliminated using Reb blood cell lysing buffer (BD Biosciences) for 15-20 min. Cells were then stained with the following mix of antibodies: PE-Cy7 rat anti-mouse CD45 (BD #552848), PE-Cy7 rat anti-mouse TER-119 (BD #557853), PE rat anti-mouse Ly6A (Sca-1; BD #553108), BV421 rat anti-mouse CD34 (BD #562608), Alexa Fluor 700 rat anti-mouse α7-integrin (Invitrogen #MA5-23607). Using a FACS Aria II cytometer, CD45^neg^, TER-119^neg^, Sca1^neg^, CD34^pos^ and α7 integrin^pos^ cells were collected.

### Primary myoblast culture and recombinant virus transduction

WT and *Eya3^-/-^Eya4^flox/flox^;Pax7^+/+^*myoblasts were grown in Growth Medium (GM; DMEM/F12 (Gibco), 20% fetal calf serum (Eurobio), 2% Ultroser G (Sartorius), 1% Anti/Anti (Penicillin-Streptomycin (10,000 U/mL), 100X Amphotericin B (ThermoFisher)), 2 mM L-glutamine (ThermoFisher)) and plated on Matrigel (Corning)-coated Petri dishes. When cells reached 80% confluency, they were trypsinized, centrifuged (400 *g*) and re-seeded at 1/3 decreased confluence for amplification. To induce the excision of the floxed *Eya4* allele in myoblasts, *Eya3^-/-^Eya4^flox/flox^* myoblasts were plated at 10,000 cells/cm^2^ concentration on Matrigel-coated plates in growth medium and transduced twice with Ad-mCherry or Ad-Cre-mCherry adenoviruses (200 MOI, Vector Biolabs). They were then FACS-sorted to get pure Adeno-positive cell populations before amplification: *Eya3^-/-^;Eya4^flox/flox^;Ad-mCherry* (*Eya3^-/-^*cells ∼ Che cells) and *Eya3^-/-^Eya4^flox/flox^;Ad-Cre-mCherry* (*Eya3^-/-^Eya4^SC/SC^* ∼ CRE cells). To study myogenic proliferation, myoblasts were plated at 20,000 cells/cm^2^ concentration on Matrigel-coated circular coverslips. To induce myogenic differentiation and fusion, myoblasts were plated at 45,000 cells/cm^2^ concentration on Matrigel-coated plates in growth medium. Once adherent, cells were incubated in Differentiation medium (DM; DMEM 1g/L glucose (Gibco), 2% horse serum, 1% Anti/Anti) for up to 3-4 days.

### Immunocytochemistry for cell culture

Muscle cells were fixed for 10 min with 4% PFA, washed several times in 1X PBS and incubated with 5% HS/0.5% Triton/0.1M glycine/1X PBS for 20 min at RT. Cells were incubated with primary antibodies during 1 hour at RT, followed by several 1X PBS washes and 1-hour incubation with the secondary antibodies. Nuclei were stained with Hoechst. Primary and secondary antibodies are listed in (Table S2). For proliferation assay, cells were incubated with EdU for 2 hours before fixing them with 4% PFA for 10 min at RT. EdU detection was performed using Click-iT EdU Alexa Fluor 647 kit, according to manufacturer’s instructions (Life Technologies). Then cells were incubated with primary and secondary antibodies as previously described.

### Image acquisition and morphometric analysis

Images were acquired on either an upright fluorescent microscope (Olympus BX63) equipped with an ORCA-Flash 4.0 LT Hamamatsu camera using MetaMorph 7 software, or an inverted confocal spinning disk microscope (IXplore Spinning IX83) equipped with a Hamamatsu sCMOS Orca Flash 4.0 V3 camera using CellSens Dimension software.

Fluorescence images were acquired using an Olympus BX63F microscope with 10x (UplanFL, NA 0.3) and 20x (UPLSAPO, NA 0.75) objectives, coupled with an ORCA-Flash4.0 LT camera (Hamamatsu), or a Zeiss Axiovert 200M microscope with 5x (PLANFLUAR, NA 0.25) and 20x (LD PLANNEOFLUAR, NA 0.4) objectives, couples with a CoolSnap-HQ camera (Photometrics), and Metamorph 7.7.5 software (Molecular Devices). Images were merged and edited in ImageJ. Background was reduced using brightness and contrast adjustments applied to the whole image.

Myofibre cross sectional area (CSA) was analyzed by using antibodies against embryonic myosin heavy chain (MYH3, DSHB #BF-G6) for samples at 4 days post-injury, or against laminin (sc-59854 or ab11575) for latter recovery timings; these antibodies mark newly formed myofibres and myofibre sarcolemma, respectively. All myofibres in the regenerated area were analysed.

### RNA extraction and RT-qPCR

TRIzol reagent (Ambion) was used to isolate RNA from WT, Che and CRE cells from proliferation and differentiation assays (Diff 0/1/2/3 days) according to manufacturer’s instructions. RNA was extracted using phenol:chloroform:iso-amyl alcohol (ThermoFisher) and then DNase I treated (Sigma #AMPD1). cDNA was synthesized from 500 ng of total RNA using random hexamer primers and SuperScript III Reverse transcriptase kit (Invitrogen) according to manufacturer’s instructions. RT-qPCRs were performed in 96-well plates on LightCycler 480 device and SYBR Green I kit (Roche) according to manufacturer’s instructions. Primers (Eurogentec) were designed with RealTime qPCR Assay tool (<u>Integrated DNA Technologies (idtdna.com)</u>) and Primer-BLAST (<u>Primer designing tool (nih.gov)</u>). cDNA was amplified using 40 cycles at 95°C for 15 sec, 60°C for 15 sec and 72°C for 15 sec. Gene expression levels were normalized to the averaged expression level of the housekeeping genes *cyclophilin* and *tbp* (Table S3).

## Statistical analysis

Data were analysed with GraphPad Prism 9 using unpaired non-parametric Mann Whitney U test or two-way ANOVA with Tukey’s multiple comparisons test. Results are presented as the means ± S.E.M. and are representative of at least two independent experiments. In all statistical analysis, p<0.05 (*) was considered significant; **p<0.01, ***p<0.001, ****p<0.0001.

## Bioinformatic analysis

Open-access scRNA-seq datasets (GSM4948866) (https://doi.org/10.1016/j.isci.2021.102838) derived from murine plantaris and soleus muscles (n=1), along with our published data from soleus, quadriceps, tibialis, plantaris, and a mixture of gastrocnemius+plantaris+soleus+tibialis+EDL muscles (https://doi.org/10.1038/s41467-020-18789-8) (n=4), were reanalyzed using the Seurat R package. We normalized, feature-selected, and integrated the individual data samples using the SCTransform function, followed by the FindIntegrationAnchors function. The data of muscle stem cell regeneration (GSE143437, n=10) were retrieved from <u>10.1016/j.celrep.2020.02.067</u> and reanalyzed as described above.

The transcriptome data (GSE159079) of murine muscles and *Pax7*-expressing muscle stem cells (MuSC) at developmental stages E15 and E18 were retrieved from [21] and reanalyzed using the Oligo R package. The data were combined according to gene symbols (n=11), normalized using the RMA method, and batch effects were corrected using the Limma R package. snRNA-seq data from Tibialis anterior day-3 post Notexin injury will be published elsewhere (C.Viaut et al, unpublished).

## Results

### Adult muscle regeneration in *Eya2^-/-^* mice

We first analyzed muscle regeneration in *Eya2^-/-^* adult mice 4 and 14 days following Notexin (NTX)-induced injury of the tibialis anterior (TA) muscle. As shown in Fig. 1, we observed a transient phenotype in mutant animals characterized by an increased number of smaller myofibers expressing MYH3 four days after injury (dpi). By 14 dpi, however, neither the number nor the size of the regenerated mutant myofibers differed significantly from those in the wild-type control group, and no MYH3+ myofibers were detectable. We therefore concluded that *Eya2*, which is expressed in both myogenic and non-myogenic cells of the muscle (Fig. S1A-C), is not essential for efficient adult muscle regeneration.

**Figure 1.**
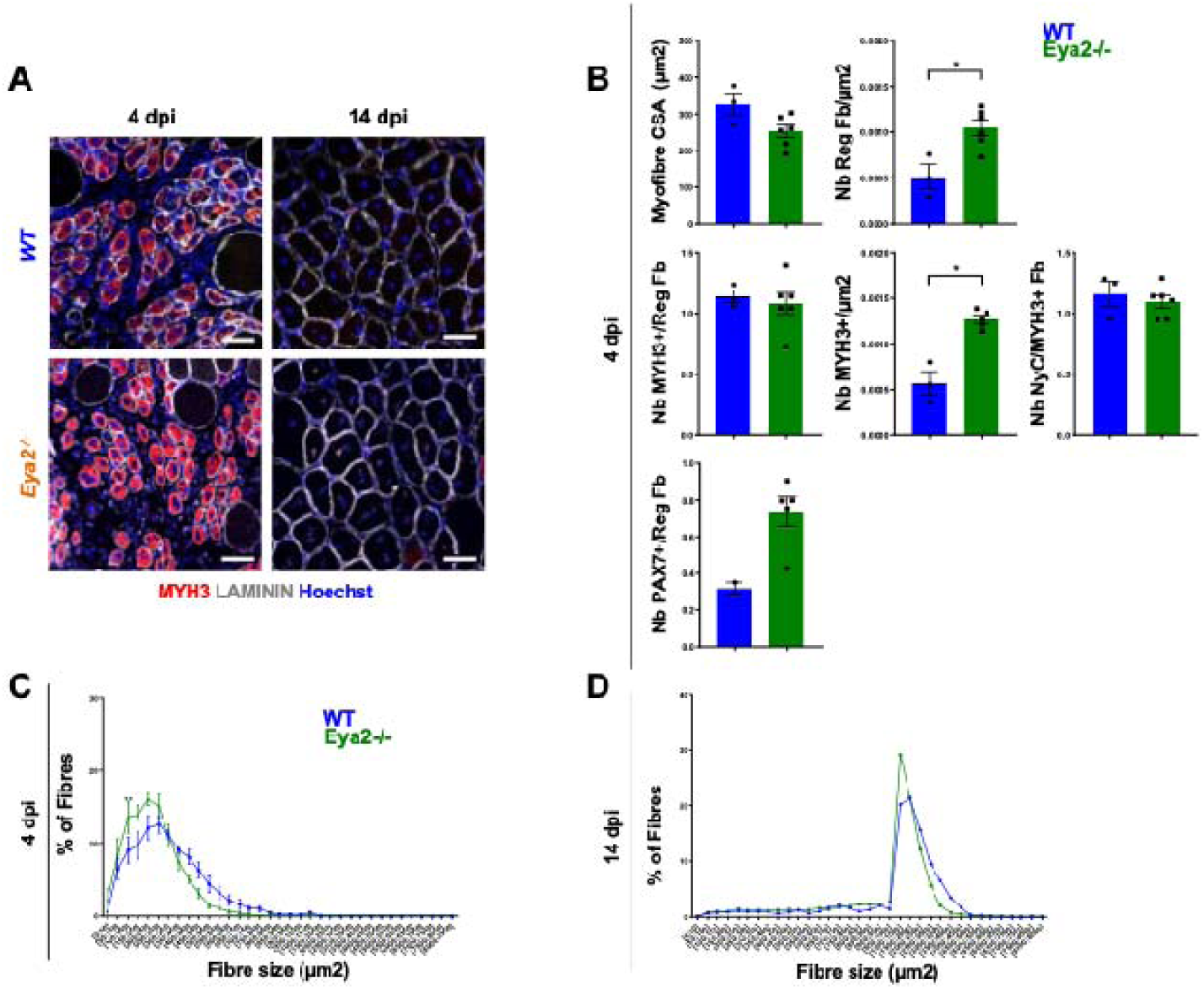
*Eya2^-/-^*mutant mice present to major muscle regeneration defects. **(A**) Immunostainings on 4 and 14 dpi of Ctrl (WT) and *Eya2^-/-^* transverse sections at the TA level for MYH3 (red), Laminin (white) and Hoechst (blue), Sb=50μm. **(B)** Quantification of MYH3, Laminin, PAX7 immunostaining for myofiber CSA per μm^2^, number of regenerated myofiber/μm^2^, number of MYH3+ myofibers on the total number of myofibers, number of regenerated MYH3+ myofiber/μm^2^, number of myonuclei present in MYH3+ myofiber section, number of PAX7+ cells by regenerated myofiber in WT and *Eya2^-/-^* regenerated TA 4 dpi. **(C)** Percentage of MYH3+ fiber size per μm^2^ in WT and *Eya2^-/-^* regenerated TA 4 dpi. **(D)** Percentage of Laminin+ fiber size per μm^2^ in WT and *Eya2^-/-^* regenerated TA 14 dpi.

### Adult muscle regeneration in *Eya3^-/-^* mice

*Eya3^-/-^* and control adult animals were treated with Tamoxifen before NTX-induced muscle injury to enable comparison with *Eya3^-/-^* ;*Eya4^sc/sc^* double mutant mice (see below). We did not detect significant impairment of muscle regeneration in *Eya3^-/-^* animals at 4- and 14-days dpi, indicating that *Eya3* alone does not play a major role in satellite cells (SCs) or other cell types contributing to adult muscle regeneration (Fig. 2).

**Figure 2.**
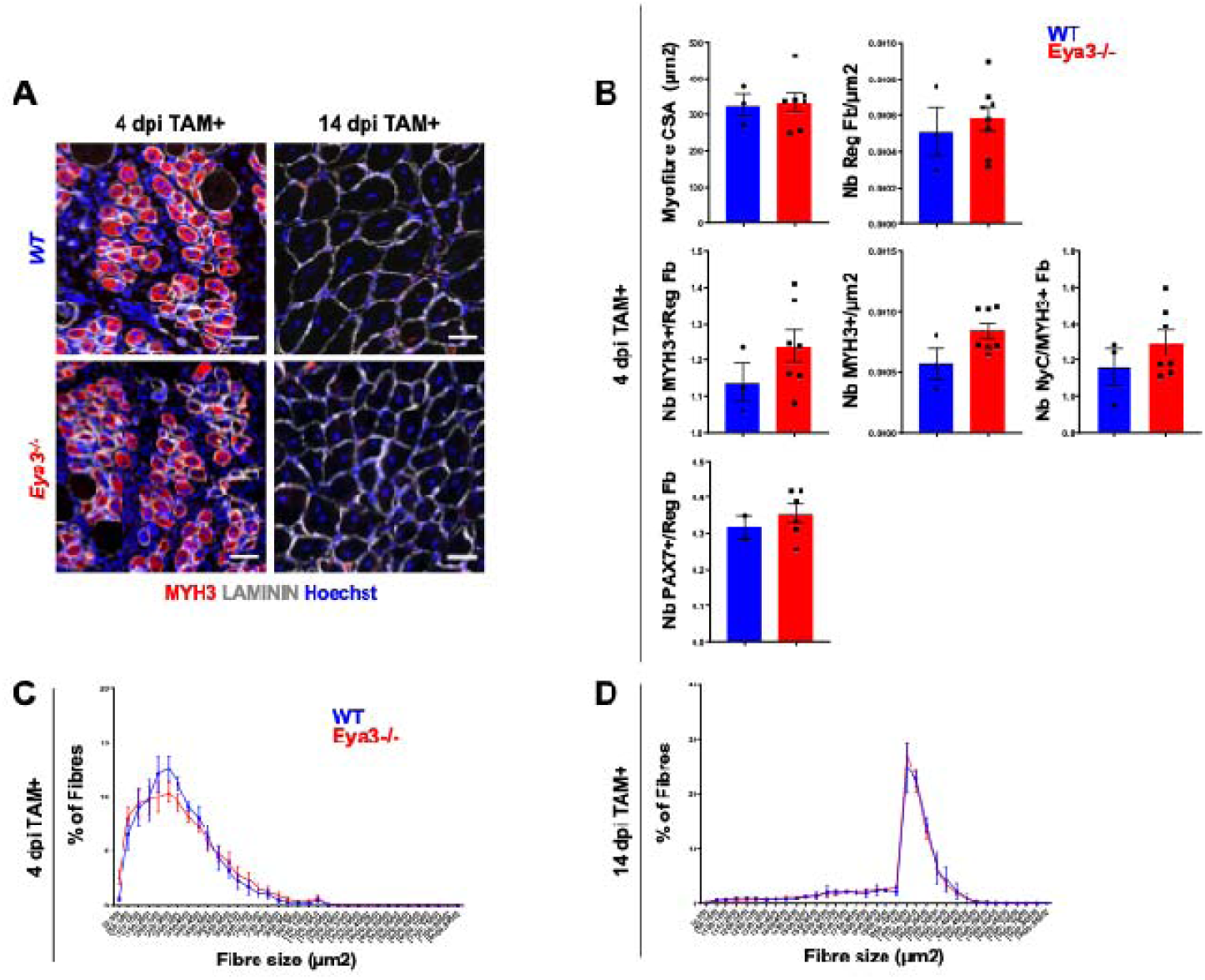
*Eya3^-/-^*mutant mice present to major muscle regeneration defects. **(A**) Immunostainings on 4 and 14 dpi of Ctrl (WT) and *Eya3^-/-^* transverse sections at the TA level for MYH3 (red), Laminin (white) and Hoechst (blue), Sb=50μm. **(B)** Quantification of MYH3, Laminin, PAX7 immunostaining for myofiber CSA in μm^2^, number of regenerated myofiber/μm^2^, number of MYH3+ myofibers on the total number of myofibers, number of regenerated MYH3+ myofiber/μm^2^, number of myonuclei present in MYH3+ myofiber section, number of PAX7+ cells by regenerated myofiber in WT and *Eya3^-/-^* regenerated TA 4 dpi. **(C)** Percentage of MYH3+ fiber size in μm^2^ in WT and *Eya3^-/-^* regenerated TA 4 dpi. **(D)** Percentage of Laminin + fiber size in μm^2^ in wt and *Eya3^-/-^* regenerated TA 14 dpi.

### Adult muscle regeneration in *Eya4^sc/sc^* mice

To investigate the role of *Eya4* in SC*, Eya4^flox/flox^* animals were crossed with *Pax7^CREert2/+^*knock-in mice [33], allowing Tamoxifen-inducible recombination of *Eya4* specifically in *Pax7*-expressing myogenic stem cells (*Eya4^sc/sc^*) (Fig. S2A-B). In parallel, *Eya3^-/-^ ;Eya4^flox/flox^;Pax7^CREert2/+^* double mutant animals were generated by crossing *Eya3^-/-^* and *Eya4^flox/flox^;Pax7^CreERT2/+^* animals. These mice were viable, fertile, and displayed no detectable phenotype prior to injury. Efficient recombination of the *Eya4^flox^* allele in SCs was confirmed by the loss of *Eya4* expression in mutant SCs compared to controls (Fig. S3A-C). Muscle regeneration after NTX injury was transiently impaired in *Eya4^sc/sc^*TA muscles (Fig. 3, Fig S4A, S5). At 4 dpi, mutant muscles displayed a reduced number of smaller MYH3+ regenerating myofibers compared with *Eya4^flox/flox^;Pax7^CreERT2/+^*controls. In addition, a transient decrease in SC number was observed during regeneration (Fig. 3B, Sup Fig. S4). However, by 14 dpi, no major differences were detected in the proportion of regenerated fast MYH4+ or slow MYH7+ myofibers (Fig. S6). These results, indicate that EYA4 in satellite cells contributes to early regenerative events, but is not required for completion of muscle regeneration.

**Figure 3.**
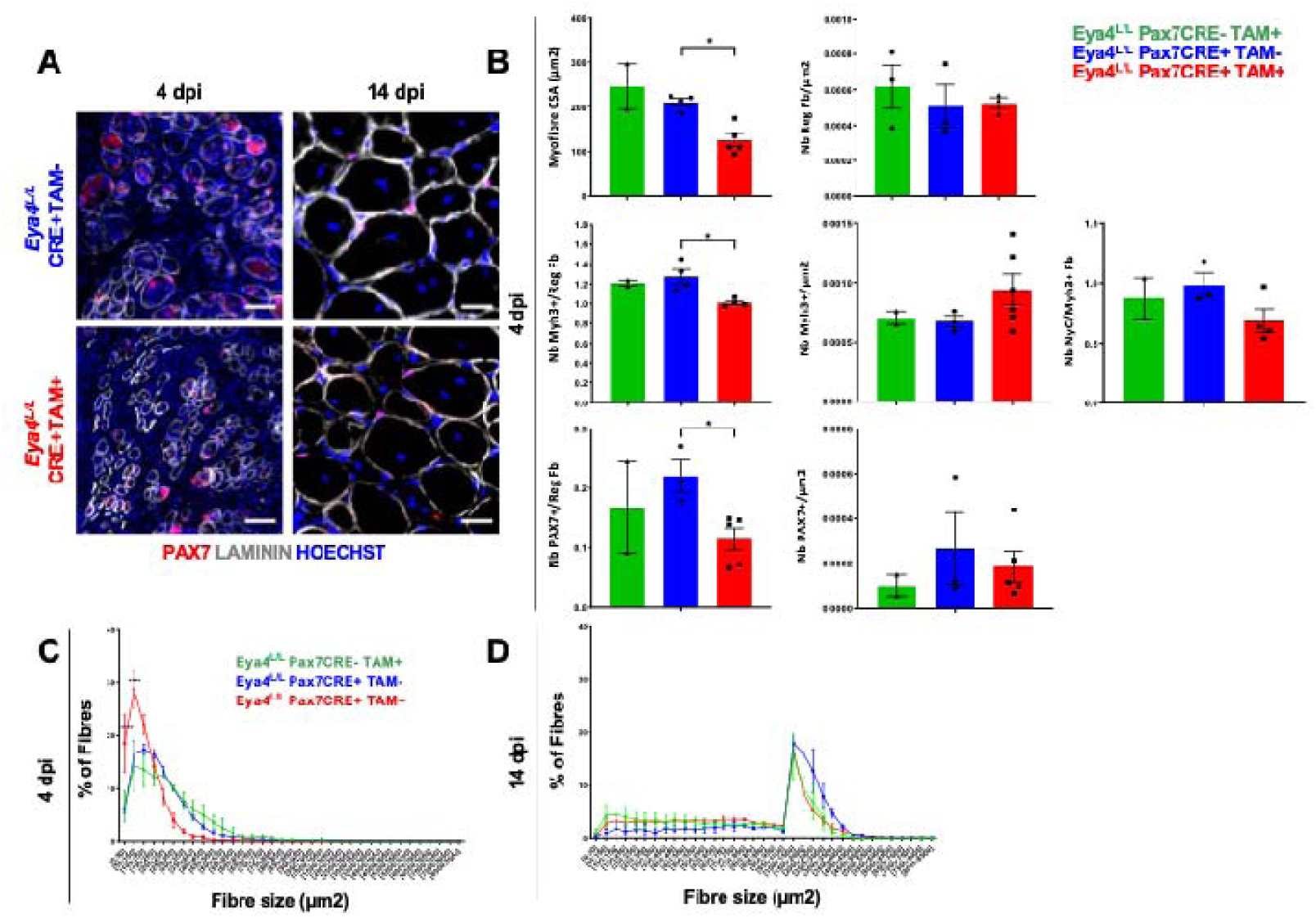
Impaired muscle regeneration in *Eya4^sc/sc^* mutant mice. **(A)** Immunostainings on 4 and 14 dpi of Ctrl (*Eya4^flox/flox^;Pax7^CreERT2/+^* without tamoxifen injection, TAM-) and *Eya4^sc/sc^* transverse sections at the TA level for PAX7 (red), Laminin (white) and Hoechst (blue), Sb=50μm. **(B)** Quantification of MYH3, Laminin, PAX7 immunostaining for myofiber CSA in μm^2^, number of regenerated myofiber/μm^2^, number of MYH3+ myofibers on the total number of myofibers, number of regenerated MYH3+ myofiber/μm^2^, number of myonuclei present in MYH3+ myofiber section, number of PAX7+ cells by regenerated myofiber, number of PAX7+ by μm^2^ in *Eya4^flox/flox^;Pax7^+/+^* with tamoxifen injection (green), *Eya4^flox/flox^Pax7^CreERT2/+^*without tamoxifen injection (blue) and in *Eya4^sc/sc^* (red) regenerated TA 4 dpi. **(C)** Percentage of MYH3+ fiber size in μm^2^ in *Eya4^flox/flox^;Pax7^+/+^* with tamoxifen injection (green), *Eya4^flox/flox^;Pax7^CreERT2/+^* without tamoxifen injection (blue) and in *Eya4^sc/sc^* (red) regenerated TA 4 dpi. **(D)** Percentage of Laminin+ fiber size per μm^2^ in *Eya4^flox/flox^;Pax7^+/+^* with tamoxifen injection (green), *Eya4^flox/flox^;Pax7^CreERT2/+^* without tamoxifen injection (blue) and in *Eya4^sc/sc^* (red) regenerated TA regenerated TA 14 dpi.

### Adult muscle regeneration in *Eya3^-/-^;Eya4^sc/sc^* mice

Because *Eya3* and *Eya4* are the two *Eya* genes most highly expressed in quiescent and regenerating SCs (Fig. S1), we tested whether these genes act redundantly during regeneration by analyzing the regenerative capacity of TA muscles from *Eya3^-/-^;Eya4^sc/sc^* mutant animals. At 4 dpi, the number of MYH3+ regenerating myofibers was increased in double mutants compared with controls (Fig. 4A-B and Sup Fig. S7). These regenerated myofibers were smaller and displayed reduced myonuclei content, consistent with defective satellite cell accretion to growing myofibers. In addition, the number of PAX7+ cells was decreased in double mutants, indicating impaired early regeneration. At 14 dpi, MYH3+ myofibers were still detected in double mutants, together with reduced CSA and decreased myonuclear content in regenerated myofibers (Fig. 4A, C-D, Fig. S7). In contrast to the decrease observed at 4 dpi, the number of SCs associated with regenerated myofibers was increased at this stage, suggesting impaired fusion of SCs into myofibers (Fig. 4C). Furthermore, while MYH4+ myofibers were readily detected in control muscles, their number was markedly reduced in double mutants, suggesting that *Eya3* and *Eya4* function not only in satellite cell fusion but also within the myofiber (Fig. S6). At 30 dpi, regenerated mutant myofibers remained smaller, displayed reduced myonuclear content, and showed an increased number of associated SCs (Fig. S7C-E). In addition, the decrease in MYH4+ myofibers was accompanied by an increase in MYH1+ myofibers (Fig. S6). Together, these results indicate that *Eya3* and *Eya4* act redundantly in SCs and are required for efficient adult muscle regeneration.

**Figure 4.**
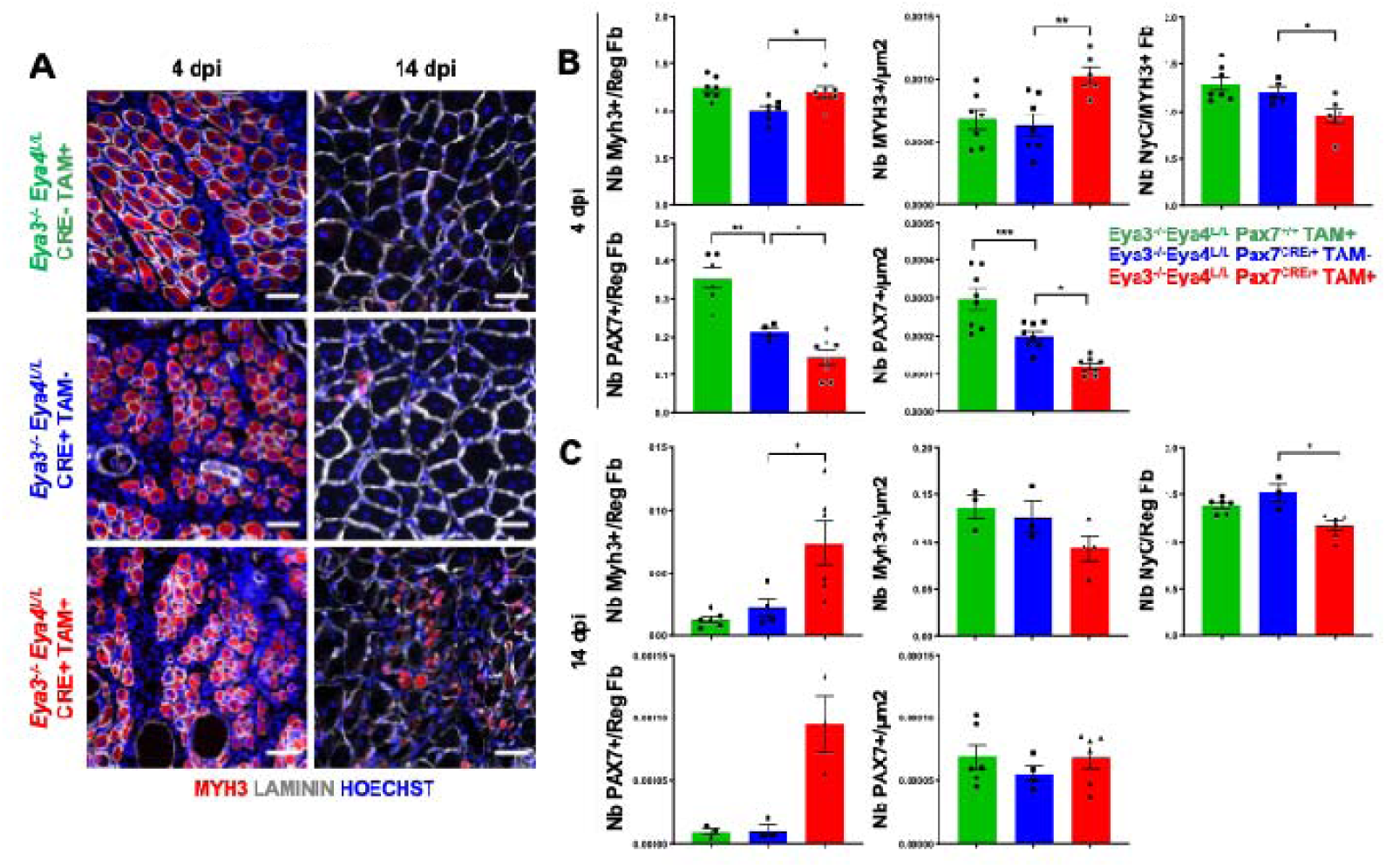
Impaired muscle regeneration in *Eya3^-/-^Eya4^sc/sc^* double mutant mice. **(A)** Immunostainings on 4 and 14 dpi of *Eya3^-/-^;Eya4^flox/flox^;Pax7^+/+^*with tamoxifen injection (green), *Eya3^-/-^;Eya4^flox/flox^;Pax7^CreERT2/+^* without tamoxifen injection, (blue, and *Eya3^-/-^ ;Eya4^sc/sc^, red,* transverse sections at the TA level for MYH3 (red), Laminin (white) and Hoechst (blue), Sb=50μm. **(B)** Quantification of MYH3, Laminin, PAX7 immunostaining for number of MYH3+ myofibers on the total number of myofibers, number of regenerated MYH3+ myofiber/μm2, number of myonuclei present in MYH3+ myofiber section, number of PAX7+ cells by regenerated myofiber, number of PAX7+ by μm^2^ *Eya3^-/-^ ;Eya4^flox/flox^;Pax7^+/+^*with tamoxifen injection (green), *Eya3^-/-^;Eya4^flox/flox^; Pax7^CreERT2/+^* without tamoxifen injection (blue), and *Eya3^-/-^;Eya4^sc/sc^* (red) regenerated TA 4 dpi and **(C)** 14 dpi.

### *Eya3^-/-^;Eya4^-/-^* satellite cells show impaired fusion properties

To characterize the cellular defects associated with *Eya3/Eya4* loss, SCs were isolated by FACS from adult *Eya3^-/-^Eya4^flox/flox^* mice And infected with adenoviruses expressing either Cherry (control) or Cherry-CRE to recombine the *Eya4^flox^*alleles, generating *Eya3^-/-^;Eya4^-/-^*SCs (Fig. S8). We first evaluated proliferation and early myogenic activation. No proliferation defects were detected in *Eya3^-/-^;Eya4^-/-^*SCs compared with control or *Eya3^-/-^*SCs, as assessed by Ki 67 staining and EdU incorporation, Fig. S8. MYOD expression was detected in control, *Eya3^-/-^* and *Eya3^-/-^;Eya4^-/-^*SC without obvious differences between genotypes (Fig. S8), indicating that early myogenic activation was preserved. Upon transfer to differentiation medium, endogenous *Eya3* and *Eya4* mRNA levels increased slightly in control cells but remained undetectable in double mutant cells (Fig. 5A). Under these conditions, *Eya3^-/-^;Eya4^-/-^* SCs displayed a marked impairment in fusion capacity (Fig. 5B, Fig. S9), accompanied by reduced activation of *Myh* genes (Fig. S10A). Expression levels of *Eya1*, *Eya2*, and *Six1* were not significantly altered in mutant SCs (Fig. S10B). To investigate the molecular basis of the fusion defect, we examined expression of fusogenic regulators. *Myomixer* expression was significantly reduced in *Eya3^-/-^;Eya4^-/-^* SCs following serum withdrawal (Fig. 5C), and decreased Myomixer protein accumulation was confirmed Western Blot analysis (Fig. 5D). Because BMP/TGFβ signaling regulates myogenic fusion [34–37] and is controlled by SIX-EYA complexes in multiple systems [38–40], we next analyzedcomponents of this pathway. While BMP4 expression showed no major change (Fig. S10C), expression of the BMP antagonists *Noggin* and *Follistatin* was strongly reduced in differentiating *Eya3^-/-^;Eya4^-/-^* SCs (Fig. 5E). This reduction was associated with increased levels of phosphorylated Smad1/5/9 proteins (Fig. 5F). Finally, analysis of published ChIPseq datasets revealed binding of SIX1/SIX4 to regulatory regions of the *Noggin*, *Follistatin* and *Myomixer* loci (Fig. S11), supporting direct transcriptioinal regulation of these genes by the SIX/EYA complex.

**Figure 5.**
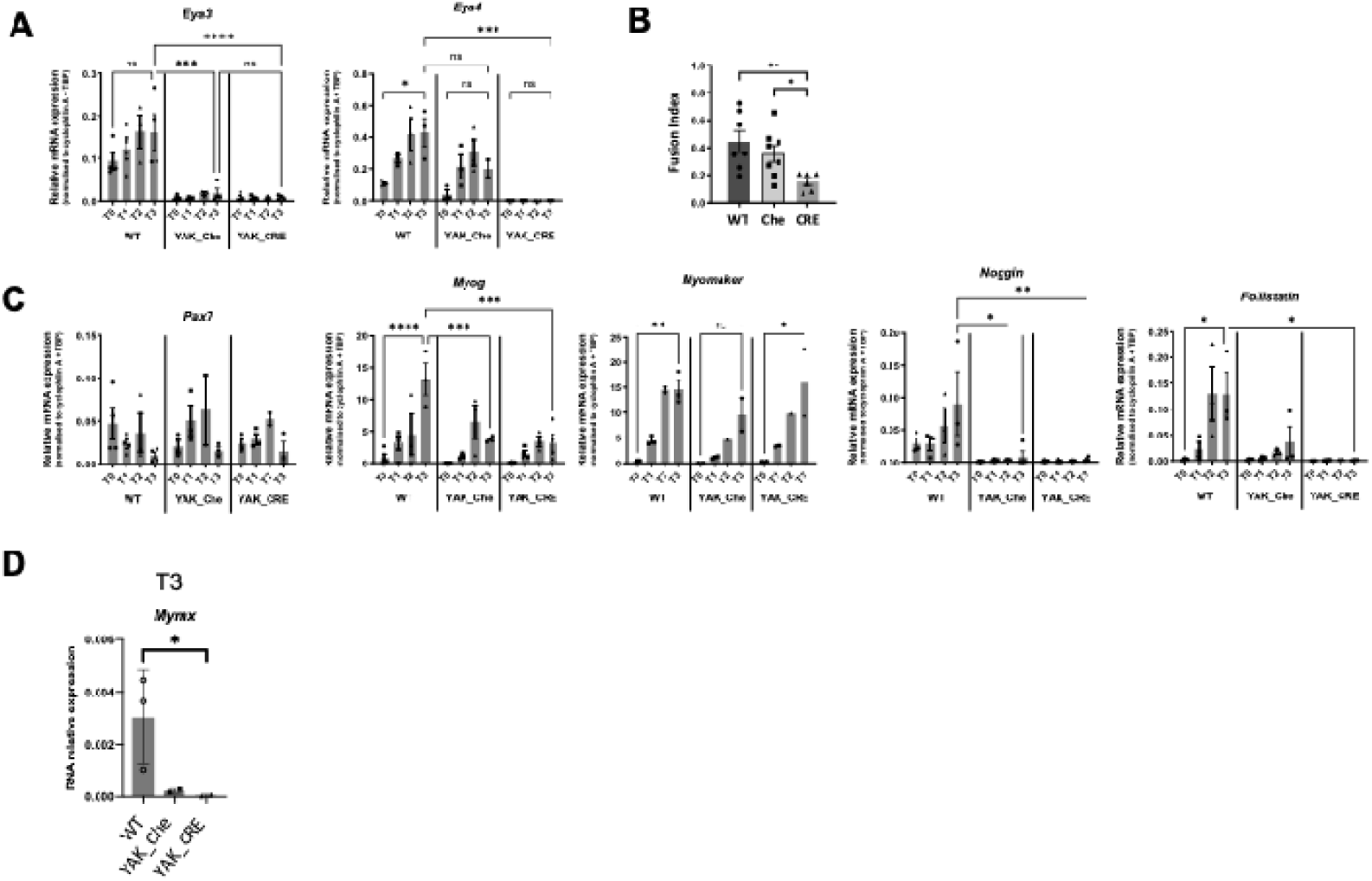
*Eya3^-/-^ Eya4^-/-^* myogenic stem cells present differentiation and fusion defects. **(A)** relative mRNA levels estimated by RT-qPCR for *Eya3* and *Eya4* at day0 (T0), day1 (T1), day2 (T2), day3 (T3) and day6 (T6) after serum withdrawal of *ex vivo* WT, *Eya3^-/-^;Eya4^flox/flox^*infected by adeno-cherry (Che) or adeno-cherry-CRE (CRE) adult myogenic stem cells. ***, p<0.001; ****, p<0.0001. **(B)** Fusion index determined by the relative number of nuclei present in cells with two or more nuclei on the total number of nuclei estimated by DAPI and WGA staining 3 days after serum withdrawal in WT, *Eya3^-/-^;Eya4^flox/flox^* infected by adeno-cherry (Che) or adeno-cherry-CRE (CRE) adult myogenic stem cells. **(C)** Same as in A for *Pax7*, *Myogenin* (Myog) and *Myomaker* gene expression. **(D)** Same as in A for and *Myomixer (Mmx)* at T0, T1, T2 and T3 (one way Anova, Kruskal-Wallis test, *,p<0.05) and for *Noggin* and *Follistatin* gene expression. At least two cell lines derived from distinct mutant mice were analyzed for each genotype.

### Eya gene function during embryonic development

Because *Eya3* and *Eya4* play major roles in adult SCs, we next examined whether these genes also regulate embryonic myogenesis. To generate constitutive *Eya4* mutants, *Eya4^flox^* animals were crossed with the ubiquitous EIIACre line to obtain *Eya4^-/+^* mice, which were subsequently crossed with the *Eya3^-/-^* animals to generate single and double mutants. The embryonic phenotypes of these mutants had not previously been established. In parallel, we further analyzed *Eya1*, *Eya2* and double *Eya1^-/-^;Eya2^-/-^*mutants, for which absence of limb musculature at E13.5 was previously reported [6]. At E18.5, *Eya3^-/-^*, *Eya4^-/-^* and double *Eya3^- /-^;Eya4^-/-^*embryos displayed no major growth defects (Fig. S12). In contrast, *Eya1^-/-^;Eya2^-/-^*fetuses exhibited severe craniofacial malformations, absence of ears, reduced body size, and bent posture, consistent with previously described phenotypes *Eya1*mutants [14]. We confirmed the absence of distal limb musculature in E18.5 *Eya1^-/-^;Eya2^-/-^* fetuses. In contrast, most distal forelimb and distal hindlimb muscles were present in *Eya3^-/-^;Eya4^-/-^*embryos, indicating that these genes are not essential for limb muscle formation (Fig. S13). Unexpectedly craniofacial musculature was severely affected in *Eya1^-/-^;Eya2^-/-^* embryos (Fig. 6A), with marked hypoplasia of extraocular muscles (EOM) and branchiomeric-derived muscles such as the masseter. In contrast, all craniofacial muscles were present in *Eya3^-/-^ ;Eya4^-/-^* embryos. Consistent with these findings, PITX2 expression was not detected in residual *Eya1^-/-^;Eya2^-/-^* EOM, whereas PITX2 was maintained in *Eya3^-/-^;Eya4^-/-^* embryos (Fig. 6B). No major hypoplasia of esophageal muscles was detected in *Eya1^-/-^;Eya2^-/-^*embryos (Fig. S13). Finally, PAX7+ myogenic stem cells were detected in craniofacial muscles of both *Eya1^-/-^;Eya2^-/-^* and *Eya3^-/-^;Eya4^-/-^* embryos at E18.5 (Fig. S13), indicating that these genes are not required for thei generation of myogenic progenitors.

**Figure 6.**
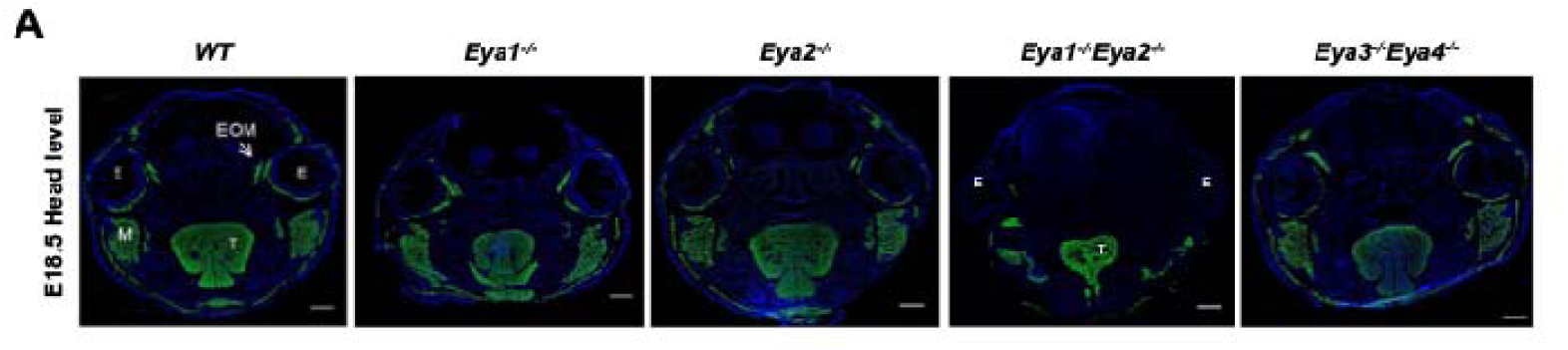
Compound *Eya* mutant fetuses show severe craniofacial muscle defects. Immunostainings on head frontal sections for E18.5 WT, *Eya1^-/-^, Eya2^-/-^, Eya1^-/-^Eya2^-/-^, Eya3^- /-^;Eya4^-/-^* fetuses at the head level for sarcomeric Myosins marked by MF20 (green), and Hoechst (blue); E: Eye, M: Masseter, EOM, extraocular muscles, T: Tongue, Sb=500μm.

## Discussion

This study defines distinct and stage-specific functions for individual EYA proteins during skeletal myogenesis. Our genetic analyses reveal that *Eya3* and *Eya4* act redundantly to support efficient adult muscle regeneration by enabling satellite cell fusion, whereas *Eya1* and *Eya2* are required during embryogenesis for proper limb and craniofacial muscle formation. Together, these findings establish a functional partitioning of *Eya* paralogs across developmental stages, linking embryonic myogenic fate acquisition to adult regenerative fusion competence.

## Comparison with *Six* mutants in hypaxial limb formation

Previous studies established that *Six1* and *Six4* control migrating hypaxial myogenesis through regulation of *Pax3* and *MRF* gene expression, and that in their absence hypaxial somitic cells fail to acquire a myogenic fate [7]. *Eya1* and *Eya2* are expressed at high levels in hypaxial somitic cells, and *Eya1^-/-^;Eya2^-/-^* double mutants were previously shown to lack myogenic cells in forelimb and hindlimb regions at E13.5 [6]. Here, we extend these observations by confirming that distal forelimbs and hindlimbs of E18.5 *Eya1^-/-^;Eya2^-/-^*mutant fetuses are devoid of myofibers and associated PAX7+ myogenic stem cells. This supports a major role for *Eya1* and *Eya2* as cofactors of SIX1 and SIX4 in activating *Pax3*-dependent programs required for hypaxial myogenic fate acquisition. In contrast, limb muscles and associated myogenic stem cells were present in E18.5 *Eya3^-/-^;Eya4^-/-^* embryos, indicating that these paralogs are not required for myogenic fate commitment in hypaxial dermomyotomal cells. Previous studies have shown that loss of *Six1* and *Six4* is associated with *Foxc1/2* upregulation, which suppresses myogenic fate acquisition [5,41]. Our observations are consistent with this regulatory framework. Together, these findings reinforce the model in which EYA1 and EYA2 act as essential cofactors of SIX proteins to drive hypaxial limb myogenesis.

## Comparison with *Six* mutants in craniofacial myogenesis

Previous studies demonstrated that *Six1* and *Six2* control craniofacial myogenesis by regulating myogenic cell fate acquisition in the cardiopharyngeal mesoderm (CPM), and that loss of these factors results in absence of CPM-derived muscles, including esophageal, trapezius and EOM [5]. In addition, reduced SIX transcriptional activity has been observed in *Eya1^-/-^* embryos using MEF3-dependent reporter assays, supporting the requirement of EYA proteins for efficient SIX transcriptional complex activity in craniofacial myogenic cells [6]. Here, we show that *Eya1^-/-^;Eya2^-/-^* double mutants present craniofacial defects including hypoplasia of branchial arch-derived muscles and EOM. Unexpectedly, esophageal musculature is preserved in these mutants, suggesting that distinct regulatory programs govern the morphogenesis of different craniofacial muscle groups [5,42], potentially reflecting compensatory activity among *Eya* paralogs or distinct lineage-specific regulatory inputs.

## Comparison with *Six* mutants in myogenic stem cells

Four *Six* genes — *Six1*, *Six2*, *Six4*, and *Six5* — are co-expressed in satellite cells and differentiated myofibers, and their contributions to adult muscle regeneration have been progressively defined. *Six1* plays a central role in SCs self-renewal and differentiation, and its absence impairs regeneration at multiple stages, including activation of *MyoD* and maintenance of the PAX7+ progenitor pool [9,10]. More broadly, SIX homeoproteins operate through feedforward transcriptional mechanisms that sustain myogenic gene expression throughout development and regeneration [12]. Consistent with this, combined loss of all four SIX proteins during embryogenesis leads to severe impairment of satellite cell self-renewal associated with loss of *Pax7* expression, demonstrating that SIX activity is required to maintain myogenic stem cell identity itself [5].

Against this background, the phenotype of *Eya3^-/-^;Eya4^-/-^*satellite cells reveals both overlapping and distinct requirements compared with SIX loss of function. EYA3/EYA4 deficiency does not eliminate the satellite cell population — PAX7+ cells are present in normal numbers in double mutant E18.5 fetuses, and adult mutant satellite cells proliferate normally and express *Myod*. These observations indicate that EYA3 and EYA4 are not essential for the satellite cell self-renewal or activation steps that depend on SIX activities. Instead, the EYA3/EYA4 requirement maps specifically to the fusion step of myogenic differentiation, downstream of *Myogenin* induction. This more restricted phenotype supports the idea that EYA proteins amplify rather than initiate SIX-dependent transcriptional programs, allowing sufficient basal activity to maintain early differentiation stages even in the absence of EYA3 and EYA4 co-activation, at least under single-injury conditions.

Our data indicate that *Eya1*, *Eya2*, *Eya3* and *Eya4* fulfill distinct functional roles in embryonic and adult myogenic stem cells, consistent with previous observations showing preferential expression of *EYA1* and *EYA2* in embryonic myogenic contexts and *EYA3* and *EYA4* in adult human satellite cells [43]. This temporal relay — from an EYA1/EYA2-dependent embryonic program to an EYA3/EYA4-dependent adult program — likely reflects the distinct transcriptional demands of embryonic myogenic fate acquisition versus adult regenerative fusion competence. The mechanism governing this transition remain unclear, but may involve changes in chromatin accessibility, shifts in SIX paralog composition, or differential abundance of EYA family members as satellite cells transition from fetal to adult identity.

We cannot exclude that inter-paralog compensation masks additional roles for EYA3 and EYA4 in satellite cell self-renewal. The persistence of *Eya1* and *Eya2* expression in *Eya3^-/-^ ;Eya4^-/-^*satellite cells may buffer early phases of the SIX-dependent program, as observed between SIX paralogs in the quadruple mutant embryo [5]. Definitive resolution of functional redundancy among Eya family members will likely require generation of satellite cell-specific quadruple *Eya* knockout models. In addition, to their role in satellite cells, *Eya* genes are expressed in differentiated myofibers, where SIX-EYA complexes regulate contractile gene expression and fiber type identity [18,44]. The reduction of MYH4+ fast fibers observed at 30 dpi in *Eya3^-/-^;Eya4^sc/sc^* regenerated muscle suggests that EYA3 and EYA4 may also contribute to fiber-autonomous specification of fast-glycolytic identity during regenerative maturation. Testing this possibility will require myofiber-specific deletion strategies to distinguish satellite cell-dependent from fiber-intrinsic roles.

## BMP signaling as a downstream effector of the EYA3/EYA4-SIX complex during regeneration

We observed a marked downregulation of *Noggin* and *Follistatin* in *Eya3^-/-^;Eya4^-/-^*differentiating satellite cells, without detectable changes in BMP4 expression. This indicates that the defect lies at the level of BMP antagonism rather than ligand production. ChIP-seq data showing SIX1 and SIX4 occupancy at the *Noggin* and *Follistatin* loci in myogenic cells further support the idea that these genes are transcriptional targets of the SIX-EYA complex. BMP signaling plays a dose-sensitive role in regulating satellite cell behavior during regeneration. BMP pathway activity supports progenitor expansion and prevents premature differentiation [45], and BMP-driven SMAD1/5/9 signaling controls the size of fetal progenitor and satellite cell pools during development [37]. Conversely, excessive BMP activity suppresses differentiation and impairs regeneration, as illustrated by defective repair in inhibitor of differentiation knockout models [46]. The balance between BMP ligands and their extracellular antagonists is therefore a critical determinant of the transition from progenitor expansion to differentiation. Follistatin regulates BMP-7 activity during embryonic muscle growth [47], Noggin controls satellite cell myogenic potential downstream of Wnt/β-catenin [48], and both are required for appropriate BMP attenuation during postnatal and regenerative myogenesis [36,49].

SIX homeoproteins have been shown to modulate BMP/TGFβ signaling in multiple developmental contexts, including kidney branching morphogenesis and craniofacial development [38–40], suggesting that the capacity of the SIX-EYA axis to tune BMP pathway output is a conserved regulatory strategy redeployed across tissues. In adult satellite cells, we establish that EYA3 and EYA4 are the critical cofactors through which SIX proteins maintain sufficient BMP antagonism to permit the transition from proliferating progenitors to fusion-competent myoblasts.

## EYA3 and EYA4 control myogenic fusion through coordinated regulation of *Myomixer* and BMP antagonists

Vertebrate myoblast fusion is a tightly orchestrated, multi-step process requiring sequential coordination of transcriptional control, cytoskeletal remodeling, and membrane merger. The transcription factor SRF (Serum Response Factor) plays a pivotal upstream role in this process by sensing the balance between monomeric G-actin and filamentous F-actin through its co-activator MAL/MRTF-A [50]. Downstream, the small GTPase RhoA must be precisely regulated — its transient activation sustains early myogenic commitment [51]. In addition to transcriptional regulation, EYA proteins have been implicated in actin cytoskeleton control through Rac and Cdc42 signaling pathways [52]. Given the established role of actin remodeling during myoblast fusion [50], defects cytoskeletal dynamics may also contribute to the impaired fusion observed in *Eya* mutant cells. The final membrane bilayer merger is then executed by the two vertebrate-specific fusogens: Myomaker, a seven-pass transmembrane protein promoting hemifusion of the outer leaflets, and Myomixer (Myomerger/Minion), a micropeptide that induces positive membrane curvature to drive fusion pore formation and expansion. Genetic loss of either fusogen completely abolishes myoblast fusion during embryogenesis and muscle regeneration, establishing this two-component machinery as both necessary and sufficient for fusogenic competence [30,53–57]. The BMP/TGFβ signaling environment further shapes fusion efficiency: TGFβ signaling acts as a molecular brake on myoblast fusion [34,35], and BMP pathway hyperactivation during differentiation is expected to have similar inhibitory consequences.

We observed a strong impairment of fusion ability in *Eya3^-/-^;Eya4^-/-^*satellite cells despite normal proliferation, *Myod* expression, and *Myogenin* induction. This fusion blockade is associated with two converging molecular defects: a failure to upregulate *Myomixer*, and a failure to suppress BMP signaling through loss of *Noggin* and *Follistatin* expression. These defects are not independent — excessive BMP-SMAD activity during differentiation is likely to further repress the pro-fusion gene program, creating a reinforcing loop in which loss of EYA-dependent BMP antagonism compounds the direct transcriptional deficit at the *Myomixer* locus. Consistent with this, SIX1 and SIX4 ChIP-seq data reveal binding peaks at the *Myomixer* locus in myogenic cells whose expression is also reduced in differentiating *Six1Six4* mutant myogenic cells [21], placing this fusogen under direct SIX-EYA transcriptional control alongside the BMP antagonists.

*Myomaker* and *Myomixer* expression is transcriptionally induced downstream of MYOGENIN.

Notably, *Myomaker* expression was not significantly altered in double mutant cells, indicating that EYA3/EYA4 selectively regulate *Myomixer* among the two fusogens. This underscores that the SIX-EYA complex acts at a defined node within the differentiation program rather than globally suppressing myogenic commitment. *In vivo*, accumulation of PAX7+ satellite cells together with reduced myonuclear content and persistent fiber size deficits at 14 and 30 dpi is consistent with a cell-autonomous defect in fusion rather than impaired activation or lineage commitment. Taken together, these observations support a model in which the SIX-EYA3/EYA4 transcriptional complex promotes fusion competence by coordinating *Myomixer* expression with maintenance of an appropriate BMP antagonist environment.

## Conclusions

In this study, we dissected the individual and combined contributions of the four mammalian EYA proteins to skeletal myogenesis across developmental time. Our results reveal a clear functional dichotomy within the EYA family: EYA1 and EYA2 are the dominant SIX cofactors in embryonic myogenic progenitors, where their combined loss abolishes hypaxial limb myogenesis and impairs craniofacial muscle development. In contrast, esophageal musculature remains preserved, suggesting the existence of distinct upstream regulatory cascades governing different craniofacial muscle groups [58]. EYA3 and EYA4, by comparison, are dispensable for embryonic myogenic fate acquisition but act redundantly in adult satellite cells to support efficient muscle regeneration.

At the adult stage, EYA4 alone contributes to early regenerative dynamics, and its satellite cell-specific loss produces a transient deficit that resolves over time. However, combined loss of EYA3 and EYA4 in satellite cells results in a persistent regenerative failure characterized by impaired myofiber growth, reduced myonuclear accretion, and fiber-type remodeling. The cellular basis of this failure is a severe impairment in myogenic fusion, associated with downregulation of *Myomixer*, *Noggin*, and *Follistatin* in differentiating double mutant cells that may increase BMP signaling. The presence of SIX1 and SIX4 binding sites at all three loci supports a model that the SIX-EYA transcriptional complex promotes a pro-fusion gene program by maintaining BMP pathway inhibition during myogenic differentiation.

Together, these findings establish EYA proteins as stage-specific and paralog-selective regulators of the SIX transcriptional complexes across the full arc of skeletal muscle development, and identify the EYA3/EYA4-SIX axis as a critical determinant of myogenic fusion competence of adult satellite cells. More broadly, these results provide a framework for understanding regenerative failure in pathological contexts where satellite cell differentiation is impaired.

## Declarations

### Ethics approval

All animal experiments were conducted in accordance with the European guidelines for the care and use of laboratory animals and were approved by the institutional ethic committee and FrenchMinistry of Research (number A751402).

Animal experimentation adhered strictly to the institutional guidelines for care and use of laboratory animals as outlined in the European Convention STE 123 and the French National Charter on the Ethics of Animal Experimentation (n°44083-2022121514506347, n°2024022718312758 and n°201809041512432, agreement n°C75-14-02). All animal procedures received approval from the French Ethical Committee of Animal Experiments CEEA - 034 and were performed in Institut Cochin animal core facility (Agreement A751402).

### Consent for publication

All authors have read and approved the final manuscript.

### Competing interests

The authors declare no competing interests.

### Funding

C.V is supported by the Agence Nationale pour la Recherche (ANR). Financial support was provided by the AFM (AFM 17406) the Institut National de la Santé et de la Recherche Médicale (INSERM), the Centre National de la Recherche Scientifique (CNRS), and the ANR (ANR16-CE-14-0002-01 BMP-MYOSTEM) to PM, and by the ANR (ANR16-CE-14-0002-01 BMP-MYOSTEM) to H.A.

### Author Contributions

Designed experiments: C.V, P.M, H.A. Performed experiments : C.V., M.W., I.P., E.J., S.B. Bioinformatics; I.P, E.J. Interpreted the data : C.V., S.B., M.W., A.S., P.M. Wrote the manuscript : P.M with input from C.V. All authors revised the manuscript. Funding Acquisition, P.M., H.A.

## Supporting information

Viaut et al, Sup data

## Acknowledgements

We thank Vladimir Ugorets and Sabrina Pichon for helpful critical reading of the manuscript, Nathalie Didier for helpful discussion concerning myogenic stem cells FACS isolation, Murielle Andrieu of the Cochin Cybio facility, Thomas Guilbert of the Cochin IMAG’IC facility, Rémi Pierre and Marcio Do-Cruzeiro of the Cochin MOUST’IC core facility and Antoine Guéraud of the Institute Cochin for their technical assistance.

## Abbreviations

Eya: eyes absent
Six: sine oculis homeobox
BMP: bone morphogenetic protein
MEF3: muscle enhancer factor 3
iPSC: induced pluripotent stem cell
SC: satellite cell
MYH: Myosin heavy chain

## Notes

### Competing Interest Statement

The authors have declared no competing interest.

