## Supplementary material for "EYA1/EYA2 and EYA3/EYA4 act as stage-specific SIX cofactors in embryonic and adult regenerative skeletal myogenesis": Viaut et al, Sup data

Supplementary Materials include : 13 supplementary figures and 3 supplementary tables.

**Legends of supplementary figures**

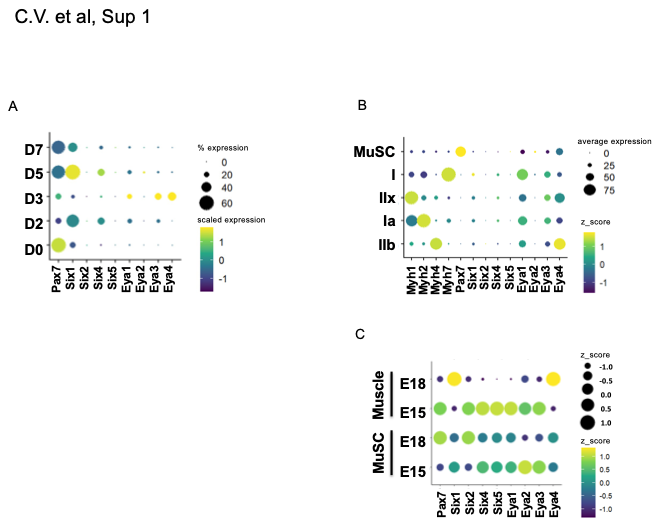
**Sup. Figure 1. Scatter plots showing the expression of *Pax7*, *Six*, *Eya* and *Myh* genes. (A)** Expression of indicated genes in adult myogenic stem cells during quiescence (D0), 2 (D2), 5 (D5) or 7 (D7) days after muscle injury, data from [59], and 3 (D3) days after Notexin injury. **(B)** Expression of indicated genes in nuclei of adult muscle stem cells (MuSC) or in IIb(Myh4+), IIa (Myh2+), IIx (Myh1+) and I (Myh7+) myonuclei, data from [60]. (**C)** Expression of indicated genes in PAX7+ myogenic cells in E15 (Pax7E15) and E18 (Pax7E18) back muscles and from back muscles at E15 or E18, data from [21]. The intensity of the expression, and the quantity of positive cells /nuclei are indicated.

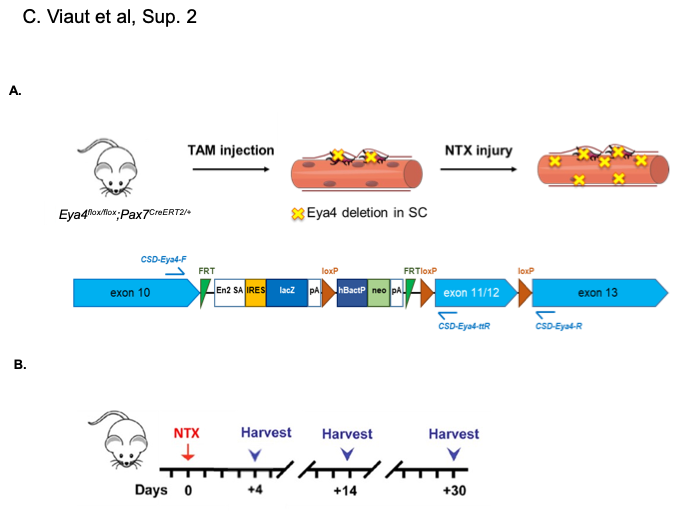
**Sup. Figure 2. *In vivo* muscle regeneration in *Eya4^flox^* animals. (**A) Up, three days after TMX treatment, TA muscles of control and *Eya4^flox/flox^* mice were injured by a single NTX injection and analyzed at various times during the muscle regeneration process. Down, deletion of *Eya4* in *Pax7+* myogenic stem cells leads to regenerated myofibers with myonuclei and associated SC mutant for *Eya4*. (B) KOMP *Eya4* recombined allele used in this study, deleting exons 11 and 12 after CRE recombination.

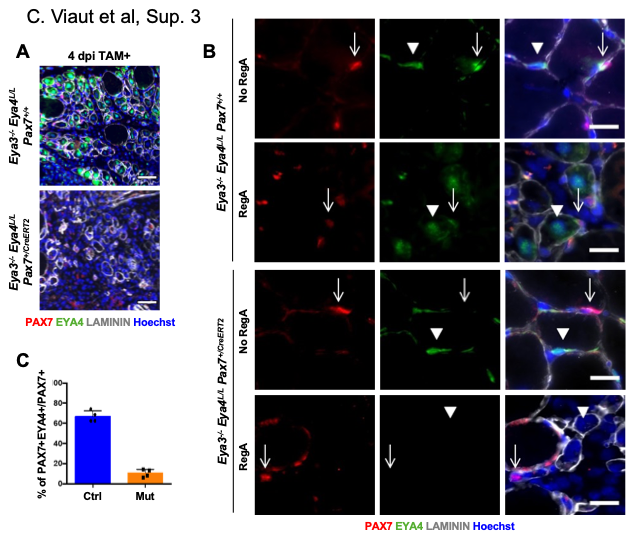
**Sup. Figure 3. EYA4 is detected in adult SC, but no more in *Eya4^sc/sc^*. (A)** Immunostainings on 4dpi of Ctrl (*Eya3^-/-^;Eya4^flox/flox^;Pax7^+/+^* and *Eya3^-/-^;Eya4^flox/flox^;Pax7^CreERT2/+^* with tamoxifen injection) transverse sections at the TA level for PAX7 (red), EYA4 (green), Laminin (white) and Hoechst (blue) Sb=50μm. **(B)** Immunostainings on 4dpi (RegA) or contralateral (no RegA) of *Eya3^-/-^;Eya4^flox/flox^;Pax7^+/+^* (up panel) and *Eya3^-/-^;Eya4^flox/flox^;Pax7^CreERT2/+^* (down panel) with tamoxifen injection, transverse sections at the TA level for PAX7 (red), EYA4 (green), Laminin (white) and Hoechst (blue) Sb=20μm. **(C)** % of PAX7+EYA4+ SC in control (Ctrl) and *Eya4^sc/sc^* (Mut) at the TA level 4dpi.

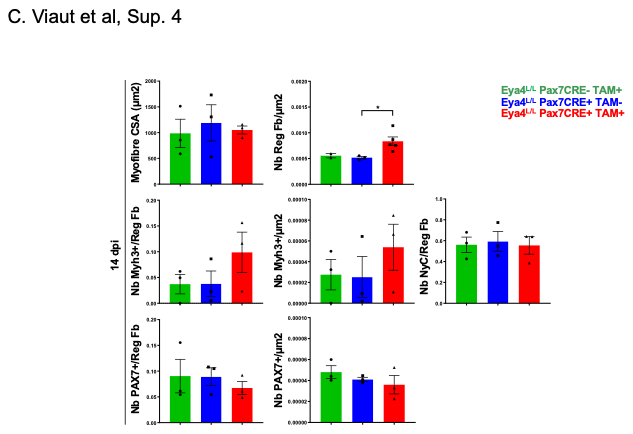
**Sup. Figure 4. Impaired muscle regeneration in *Eya4^sc/sc^* mutant mice at 14 dpi.** Quantification of MYH3, Laminin, PAX7 immunostaining for myofiber CSA in μm^2^, number of regenerated myofiber/μm^2^, number of MYH3+ myofibers on the total number of regenerated myofibers, number of regenerated MYH3+ myofiber/μm^2^, number of myonuclei present in MYH3+ myofiber section, number of PAX7+ cells by regenerated myofiber, number of PAX7+ by μm^2^ in *Eya4^flox/flox^;Pax7^+/+^* with tamoxifen injection -green, *Eya4^flox/flox^;Pax7^CreERT2/+^* without tamoxifen injection –blue- and in *Eya4^sc/sc^* –red- regenerated TA 14 dpi.

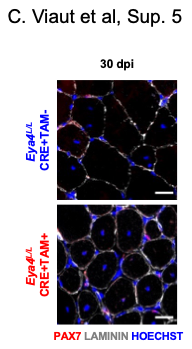
**Sup. Figure 5. Phenotype of the *Eya4^sc/sc^* mutant TA 30 dpi. (A)** Immunostainings on 30 dpi of Ctrl (*Eya4^flox/flox^;Pax7^CreERT2/+^* without tamoxifen injection) and *Eya4^sc/sc^* transverse sections at the TA level for PAX7 (red), Laminin (white) and Hoechst (blue), Sb=50μm.

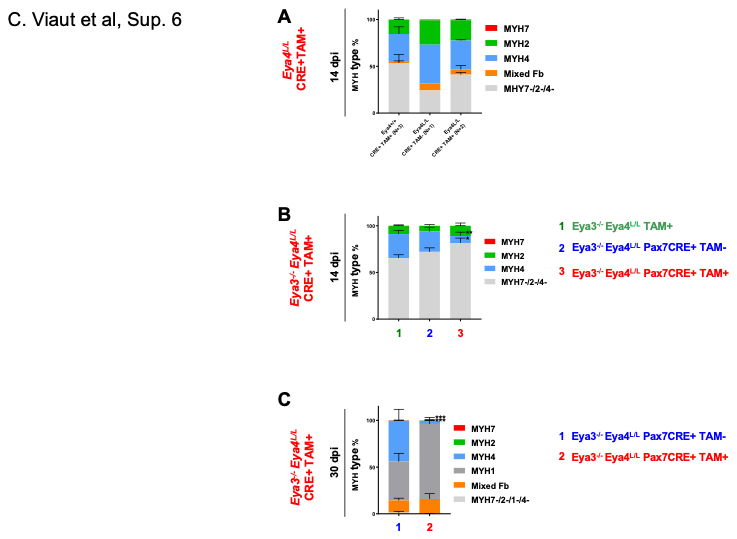
**Sup. Figure 6. Muscle fiber specialization in *Eya4^sc/sc^* and *Eya3^-/-^Eya4^sc/sc^* mutant regenerated myofibers. (A)** Percentage of MYH7+, MYH2+, MYH4+, mixed MYH2+MYH7+ and MYH2+MYH4+, and MYH7-/MYH2-/MYH4- myofibers in *Pax7^CREert2/+^*+tamoxifen (a), *Eya4^flox/flox^;Pax7^CreERT2/+^* without tamoxifen (b) and *Eya4^flox/flox^;Pax7^CreERT2/+^* with tamoxifen injection (c), 14 dpi. Three sections were analyzed by regenerated TA, the number of animals studied is indicated. **(B)** Percentage of MYH7+, MYH2+, MYH4+ and MYH7-/MYH2-/MYH4- myofibers in *Eya3^-/-^;Eya4^flox/flox^;Pax7^+/+^* with tamoxifen injection, 1-green, *Eya3^-/-^Eya4^flox/flox^Pax7^CreERT2/+^* without tamoxifen injection, 2-blue, and *Eya3^-/-^Eya4^sc/sc^,*3- red regenerated TA 14 dpi. Three sections were analyzed by regenerated TA, three or more animals were studied. **, p<0.01. **(C)** Percentage of MYH7+, MYH2+, MYH4+, MYH1+, mixed and MYH7-/MYH2-/MYH4- myofibers in *Eya3^-/-^;Eya4^flox/flox^;Pax7^CreERT2/+^* without tamoxifen (1) and *Eya3^-/-^;Eya4^flox/flox^;Pax7^CreERT2/+^* with tamoxifen injection (2), 30 dpi. Three sections were analyzed by regenerated TA, three or more animals were studied. ***, p<0.001.

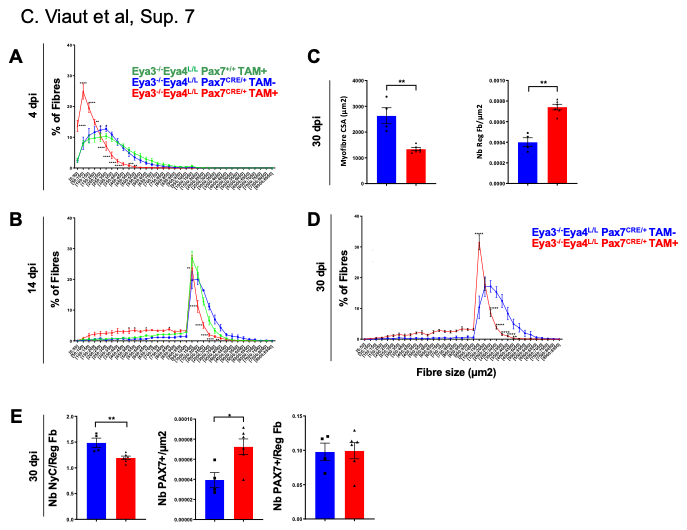
**Sup. Figure 7. Impaired muscle regeneration in *Eya3^-/-^Eya4^sc/sc^* double mutant mice, 4, 14 and 30 dpi. (A)** Percentage of MYH3+ fiber size per μm^2^ 4 dpi in *Eya3^-/-^Eya4^flox/flox^Pax7^+/+^* with tamoxifen injection -green, *Eya3^-/-^;Eya4^flox/flox^;Pax7^CreERT2/+^* without tamoxifen injection –blue- and in *Eya3^-/-^Eya4^sc/^*^sc^-red in regenerated TA. **(B)** % of Laminin + fiber size in μm^2^ 14 dpi of *Eya3^-/-^;Eya4^flox/flox^;Pax7^+/+^* with tamoxifen injection -green, *Eya3^-/-^;Eya4^flox/flox^;Pax7^CreERT2/+^* without tamoxifen injection –blue- and in *Eya3^-/-^;Eya4^sc/^*^sc^-red in regenerated TA 14 dpi. **(C)** Quantification of Laminin and Hoescht immunostaining for CSA of regenerated myofibers and number of regenerated Laminin+ myofiber/μm2, in *Eya3^-/-^;Eya4^flox/flox^;Pax7^CreERT2/+^* without tamoxifen injection, blue, and *Eya3^-/-^;Eya4^sc/sc^,* red*,* regenerated TA 30 dpi and **(D)** Percentage of Laminin + fiber size per μm^2^ of *Eya3^-/-^;Eya4^flox/flox^;Pax7^CreERT2/+^* without tamoxifen injection –blue- and in *Eya3^-/-^;Eya4^sc/^*^sc^-red in regenerated TA 30 dpi. **(E)** number of myonuclei present in Laminin+ myofiber section, number of PAX7+ per μm^2^ and number of PAX7+ cells per regenerated myofiber, in *Eya3^-/-^;Eya4^flox/flox^;Pax7^CreERT2/+^* without tamoxifen injection –blue- and in *Eya3^-/-^;Eya4^sc/^*^sc^-red in regenerated TA 30 dpi. *, p<0.05; **, p<0.01; ***, p<0.001; ****, p<0.0001.

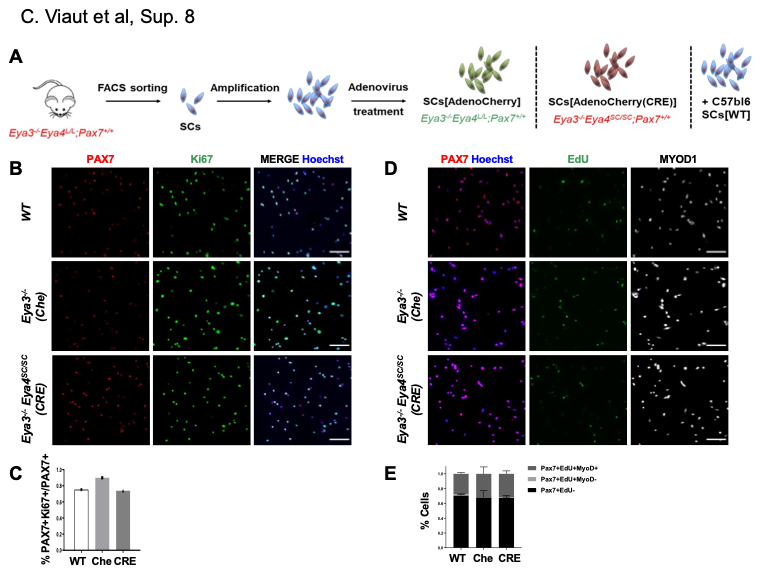
**Sup. Figure 8. *Eya3^-/-^ Eya4^-/-^* myogenic stem cells present no proliferation defects when cultured *ex vivo*. (A)** General diagram explaining how mutant myogenic stem cells are obtained after FACS sorting, amplification in culture and infection with recombinant adenovirus. **(B)** WT, *Eya3^-/-^;Eya4^flox/flox^* (Che) and *Eya3^-/-^;Eya4^-/-^* (Cre) were amplified *ex vivo* fixed and hybridized with PAX7 (red) and Ki67 (green) antibodies. Hoechst revealed the nuclei. **(C)** Histogram of the quantification of the percentage of Ki67+PAX7+ cells on total PAX7+ cells. **(D)** WT, *Eya3^-/-^;Eya4^flox/flox^* (Che) and *Eya3^-/-^;Eya4^-/-^* (Cre) were amplified *ex vivo* and treated 3 hours with EdU, fixed and hybridized with PAX7 (red), EdU (green) and MYOD antibodies. Hoechst revealed the nuclei. **(E)** Histogram of the quantification of the percentage of PAX7+EdU+MYOD+ (dark grey) PAX7+EdU+MYOD- (light grey) and PAX7+EdU- (black) in WT, *Eya3^-/-^;Eya4^flox/flox^* (Che) and *Eya3^-/-^;Eya4^-/-^* (CRE) cells.

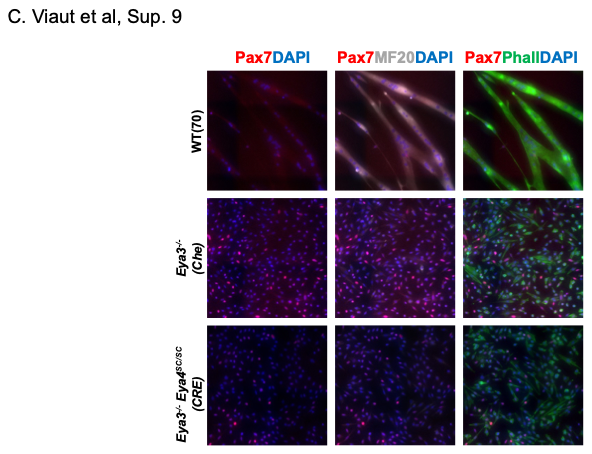
**Sup. Figure 9. *Eya3^-/-^ Eya4^-/-^* myogenic stem cells present fusion defects when differentiated *ex vivo*.** WT, *Eya3^-/-^;Eya4^flox/flox^* (Che) and *Eya3^-/-^;Eya4^-/-^* (Cre) were amplified *ex vivo* fixed and hybridized with PAX7 (red) and MF20 (grey) antibodies and Phalloidin (Phall, green). DAPI revealed the nuclei.

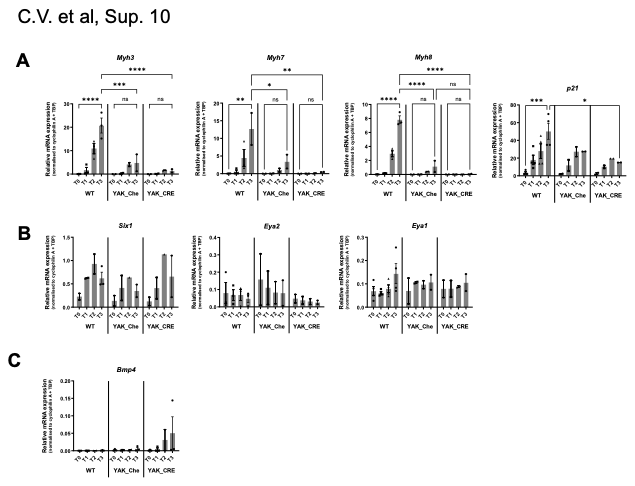
**Sup. Figure 10. *Eya3^-/-^ Eya4^-/-^* myogenic stem cells present differentiation defects when cultured *ex vivo*. (A-C)** relative mRNA levels estimated by RT-qPCR for (**A**) *Myh3, Myh7, Myh8*, (**B**) *Six1, Eya1, Eya2* and (**C**) *BMP4* at day0 (T0), day1 (T1), day2 (T2), day3 (T3) and day6 (T6) after serum withdrawal of *ex vivo* WT, *Eya3^-/-^;Eya4^flox/flox^* infected by adeno-cherry (Che) or adeno-cherry-CRE (CRE) adult myogenic stem cells. *,p<0.05; **,p<0.01; ***,p<0.001; ****, p<0.0001.

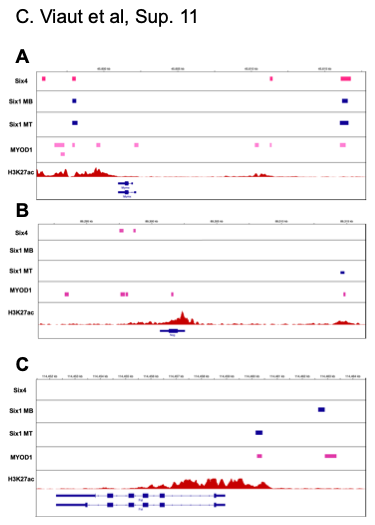
**Sup. Figure 11. Binding of SIX homeoproteins at the *Noggin*, *Myomixer* and *Follistatin* loci.** SIX1 [61], SIX4 [8], MYOD1, H3K27ac ChIP-seq experiments at the *Myomixer* (Mymx, A), *Noggin* (Nog, B) and *Follistatin* (Fst, C) mouse loci in muscle cells, and showing peaks of SIX1 and/or SIX4 binding in opened DNA regions (H3K27ac peaks) that may correspond to promoter and enhancer elements. MB, myoblast. MT, myotube.

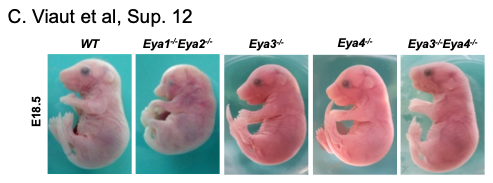
**Sup. Figure 12. Representative images of WT, *Eya1^-/-^Eya2^-/-^; Eya3^-/-^, Eya4^-/-^,* and *Eya3^-/-^Eya4^-/-^E18.5* fetuses.**

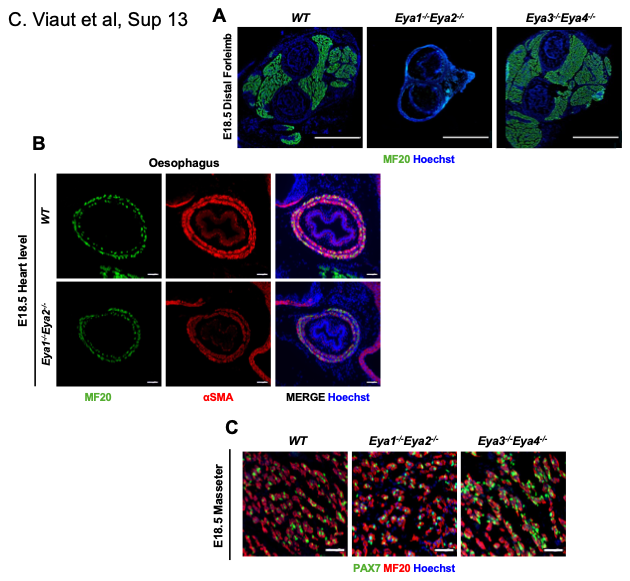
**Sup. Figure 13. PAX7+ myogenic stem cells are present in compound E18.5 *Eya* mutant fetuses. (A)** Immunostaining with MF20 antibodies revealing all sarcomeric Myosin heavy chains and Hoechst at the distal forelimb of E18.5 WT, *Eya1^-/-^;Eya2^-/-^*and *Eya3^-/-^;Eya4^-/-^* fetuses. **(B)** Immunostaining with MF20 (green) and αSMA (red) antibodies and Hoechst at the heart level to reveal skeletal and smooth muscles of the esophagus of E18.5 WT and *Eya1^-/-^;Eya2^-/-^*fetuses. **(C)** Immunostaining with PAX7 (green), MF20 (red) antibodies and Hoechst at the masseter level of E18.5 *Eya1^-/-^;Eya2^-/-^*and *Eya3^-/-^;Eya4^-/-^* fetuses showing the presence of PAX7+ cells associated with myofibers.

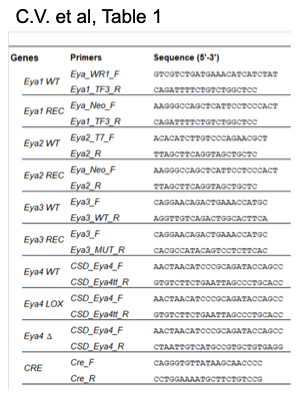
**Sup. Table 1. List and sequences of the DNA primers used for mice genotyping**

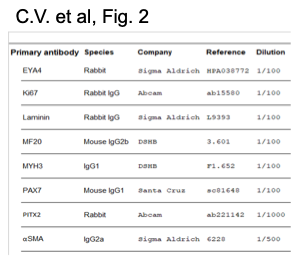
**Sup. Table 2. List of antibodies used in this study.**

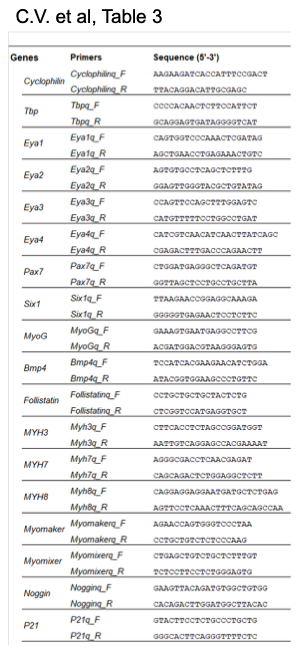
**Sup. Table 3. List and sequences of the DNA primers used to perform qPCR experiments.**
